# Molecular mechanism of SARS-CoV-2 components caused ARDS in murine model

**DOI:** 10.1101/2020.06.07.119032

**Authors:** Tingxuan Gu, Simin Zhao, Guoguo Jin, Mengqiu Song, Yafei Zhi, Ran Zhao, Fayang Ma, Yaqiu Zheng, Keke Wang, Hui Liu, Mingxia Xin, Wei Han, Xiang Li, Christopher D Dong, Kangdong Liu, Zigang Dong

## Abstract

COVID-19 has become a major challenge to global health, and until now, no efficient antiviral agents have been developed. The SARS-CoV-2 infection is characterized by pulmonary and systemic inflammation in severe patients, and acute respiratory distress syndrome (ARDS) caused respiratory failure contributes to most mortalities. There is an urgent need for developing effective drugs and vaccines against SARS-CoV-2 and COVID-19 caused ARDS. However, most researchers cannot perform SARS-CoV-2 related researches due to lacking P3 or P4 facility. We developed a non-infectious, highly safety, time-saving SARS-CoV-2 components induced murine model to study the SARS-CoV-2 caused ARDS and cytokine storm syndrome (CSS). We also investigated mAbs and inhibitors which potentially neutralize the pro-inflammatory phenotype of COVID-19, and found that anti-IL-1α, anti-IL-6, anti-TNFα, anti-GM-CSF mAbs, p38 inhibitor, and JAK inhibitor partially relieved CSS. Besides, anti-IL-6, anti-TNFα, anti-GM-CSF mAbs and inhibitors of p38, ERK, and MPO somewhat reduced neutrophilic alveolitis in the lung. In all, we established the murine model mimic of COVID-19, opening a biosafety and less time-consuming avenue for clarifying the mechanism of ARDS and CSS in COVID-19 and developing the therapeutic drugs.

## Introduction

In early December 2019, pneumonia of an unknown cause appeared in Wuhan. The virus strain was first isolated from patient samples on January 7, 2020, and the whole genome sequence of the virus was then obtained on January 10, 2019^1-3^. The genome of the virus was compared with SARS-CoV and MERS-CoV, and the homology was about 79% and 50% respectively. Therefore, it was named as novel coronavirus (SARS-CoV-2)^4^. COVID-19 is caused by SARS-CoV-2 infection^5^. A total of 7,410,510 confirmed cases of COVID-19 caused by SARS-CoV-2 have been reported globally as of June 14^th,^ 2020. The total number of deaths worldwide reached 418,294. Until now, no specific antiviral drugs or vaccines for SARS-CoV-2 have been developed. The epidemic of COVID-19 is threatening public health. Methods to control the virus and improve the treatments have quickly become urgent issues concerning national security across the world.

As the COVID-19 progressing, patients may develop acute respiratory distress syndrome (ARDS), cytokine storm syndrome (CSS)^6^. CSS, which showed a disorder of the immune system and a rapid increase of pro-inflammatory cytokine levels after the stimulus, has been reported to associate with the deterioration of various severe diseases^7^. In COVID-19 patients, the CSS occurred when the plasma levels of pro-inflammatory cytokines as IL-6, IL-1β, IL-2, IL-8, IL-17, G-CSF, GM-CSF, IP10, MCP1, MIP1α and TNFα were significantly increased, recruiting immune cells through a positive feedback loop, eventually forming a cytokine storm^8-10^. Therefore, clarifying the mechanism of CSS in COVID-19 patients and discovering the key cytokines contribute to developing more effective methods to block CSS, which is of great significance for the treatment of COVID-19 patients. The continued global epidemic of COVID-19 emphasizes the necessity of establishing a COVID-19 related pneumonia animal model. Currently, most animal models have used virus strains isolated from COVID-19 patients, which can simulate human pathophysiological processes, but it lacks biosecurity and can only be performed in the biosecurity P3/P4 labs^11,12^. Therefore, there is an urgent need to establish an effective animal model for the research of COVID-19 pneumonia that is non-infectious and can be performed in less well-equipped laboratories.

Here, our laboratory has successfully developed a highly biosecurity and time-saving ARDS animal model, which simulates the CSS and the pathophysiological changes happened in the COVID-19 patients by using Poly I:C and SARS-CoV-2 spike protein (Poly I:C and SP). The combination of Poly I:C and SP were used as SARS-CoV-2 mimic. We identified some neutralizing mAbs and inhibitors have some prophylactic efficacy for COVID-19. More importantly, it could have useful implications for the intervention of current and afterward coronavirus infectious diseases.

## Results

First, we inoculated SARS-CoV-2 mimic into mice lung through intratracheal injection (Figure 1a). After the administration of Poly I:C and SP, acute lung injury featuring neutrophilic inflammation and interstitial edema were observed. The lung tissue showed severe injury with Poly I:C and SP, which was more severe than Poly I:C induced only. To better understand the time course of pathologic changes, we explored the time point of lung injury. The lung injury occurred at 6 h, was most severe at 24 h, but gradually decreased at 48 h (Figure 1b). Besides, the mice suffered from extensive pleural fluid accumulation between 24 h and 48 h post-challenge. Pleural effusion was measured by using small animal MRI. Gross lung lesions were observed at 24 h after challenge, reduced at 48 h, and not apparent at 6 h (Figure 1c).

**Figure 1:**
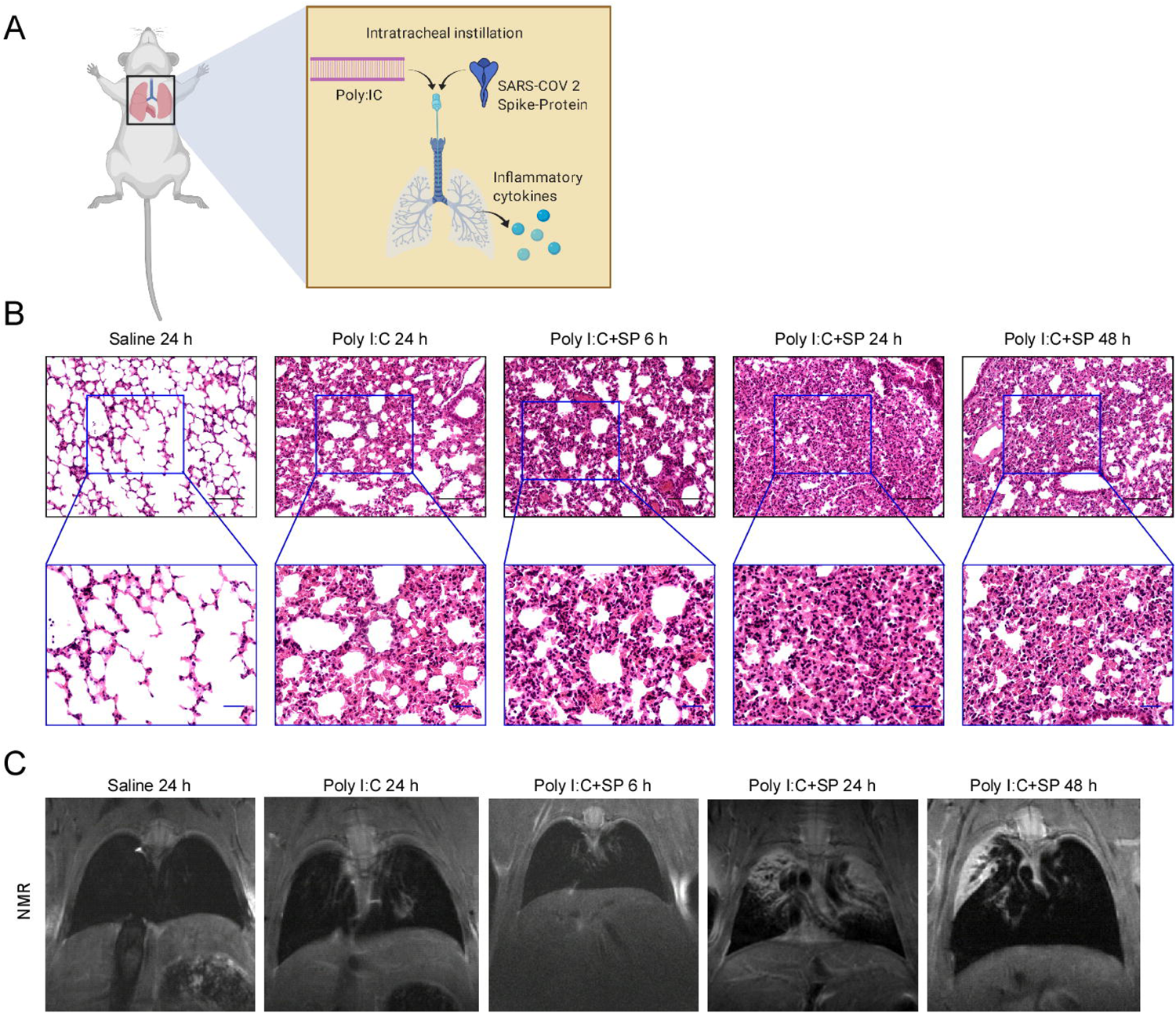
SARS-CoV-2 mimic induced acute lung injury in BALB/c mice. (a) A schematic description of Poly I:C and SARS-CoV-2 spike protein intratracheal administration in mice. (b) Histological characteristics of lung injury and interstitial pneumonia induced by Poly I:C and SARS-CoV-2 spike protein (Poly I:C + SP); 6 h, 24 h and 48 h after challenge. Control group; Saline and Poly I:C 24 h challenge. Black scale bar = 100 μm, blue scale bar = 50 μm. (c) In vivo small animal MRI documented pleural effusion induced by Poly I:C and SARS-CoV-2 spike protein; 6 h, 24 h and 48 h after challenge. Control group; Saline and Poly I:C 24 h challenge.

To understand the molecular pathology of Poly I:C and SP induced lung injury, cytokines and flow cytometric analysis were performed to determine the inflammatory molecules and related immune cells. The level of IL-1α, IL-6 and TNFα were dramatically increased in Poly I:C plus SP group as compared with Poly I:C or SP alone group. Saline and recombinant FC protein cannot stimulate the production of cytokines, Poly I:C or SP shows a higher level of IL-1α, IL-6, and TNFα (Figure 2a). Intratracheal injection of saline, SP and recombinant FC protein cannot stimulate an increase of dsDNA, an indicator of NETs (Figure 2b). Total cellularity of mice bronchoalveolar lavage (BAL) samples showed a significant increase in an SP dose-dependent manner after 24 h challenge (Figure 2c). Furthermore, investigation of cellularity in the mice BAL samples by flow cytometry. The result showed that neutrophils, but not macrophages, significantly infiltrated and migrated into the lungs. It should be noted that the number of neutrophils in the BAL at SP dose-dependent increased compared with saline (Figure 2d). For the time point study, the SARS-CoV-2 mimic led to an increase in the inflammatory cytokines IL-1α, IL-6 and TNFα at 6 h. However, TNFα and IL-6 significantly decreased at 24 h timepoint as compare with 6 h timepoint. When compared with saline, all cytokines levels were increased at 24 h timepoint (Figure 2e). We noted that there was no significant difference in the infiltration of neutrophils between those two time points (Figure 2f). Moreover, a unique mechanism of the neutrophil effector is the generation of neutrophil extracellular traps (NETs); the level of NETs can be quantified by measuring cell-free double-stranded DNA (dsDNA). In our study, the concentration of dsDNA was used as an indicator of NETs. At the 6 h and 24 h time point after challenge, there was no significant difference in the concentration of dsDNA in the mice BAL samples (Figure 2g). Furthermore, we dissociated cells from the mice lung tissue after 24 h installation of Poly I:C and SP, and maintained for 6 h and 24 h. The concentrations of IL-6 were still increasing after dissociation, and longer maintaining produced more IL-6 into the medium. This indicated that CSS happens in the lungs after installation of SARS-CoV-2, along with immune cells such as tissue-resident macrophages highly inflammatory and activated (Figure 2h).

**Figure 2:**
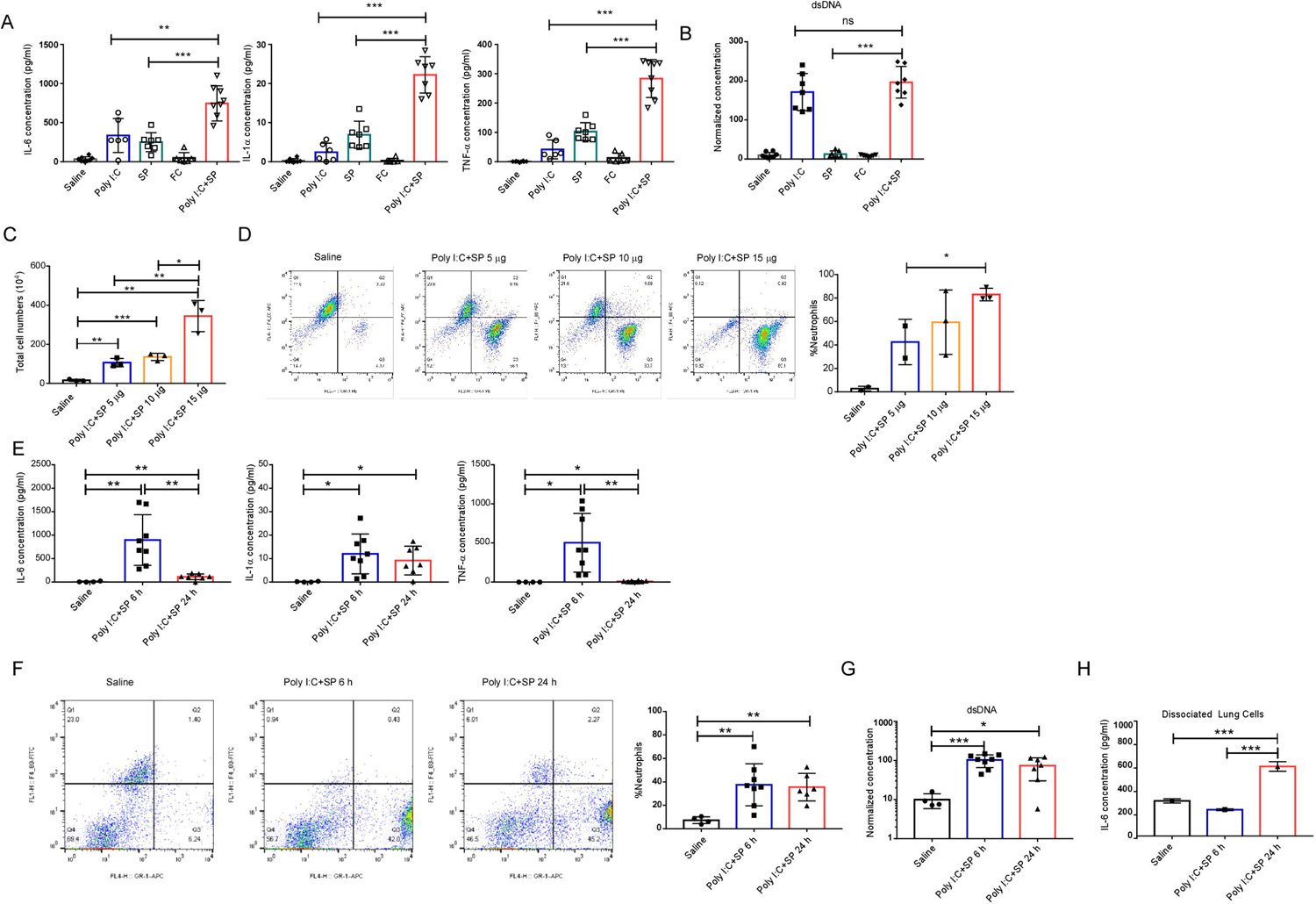
Poly I:C plus SP induced cytokines release storm in lung. (a) Production of IL-6 in the BAL from Poly I:C plus SP challenged mice, \*\**P* < 0.01, *** *P* < 0.001. (b) Levels of dsDNA in the BAL from Poly I:C plus SP challenged mice, *** *P* < 0.001. (c) Total cell count in the BAL from Poly I:C and 5 μg, 10 μg or 15 μg SARS-CoV-2 spike challenged mice, control group; Saline. * *P* < 0.05, ** *P* < 0.01, *** *P* < 0.001. 6 h or 24 h after SARS-CoV-2 mimic challenged mice. Control group; Saline (d) Cellular composition in BAL, * *P* < 0.05. (e) BAL IL-6 concentration. * *P* < 0.05, ** *P* < 0.01, *** *P* < 0.001. (f) Cellular composition in BAL, ** *P* < 0.01, *** *P* < 0.001. (g) Levels of dsDNA in the BAL. * *P* < 0.05, *** *P* < 0.001. Single cells dissociated from Poly I:C plus SP challenged mice lung (h) IL-6 concentration from cell culture medium, *** *P* < 0.001.

We hypothesized that one or more cytokines drove the CSS. To validate the hypothesis, we applied the neutralizing antibody such as anti-IL-1α, anti-IL-6 or anti-TNFα and blocking antibody anti-IL6R or anti-TNFR2 in this animal model. Neutralizing IL-6 reduced the production of cytokine IL-6 in BAL, with no effect on IL-1α and TNFα (Figure 3a). However, IL-6R blocking mAbs was unable to alter the concentrations of IL-1α, IL-6 and TNFα in BAL (Figure 3b). Anti-TNFα mAbs reduced the production of cytokine TNFα in BAL, with no effect on IL-1α and IL-6 (Figure 3c). However, TNFR2 mAbs failed to alter the production of IL-1α, IL-6 and TNFα (Figure 3d). Neutralizing mAbs targeting IL-1α directly reduced the cytokines IL-1α, and IL-6 also reduced significantly, but not TNFα (Figure 3e). In addition, concentrations of GM-CSF not increased in BAL during SARS-CoV-2 challenge, but anti-GM-CSF mAbs could reduce the production of TNFα (Figure 3f).

**Figure 3:**
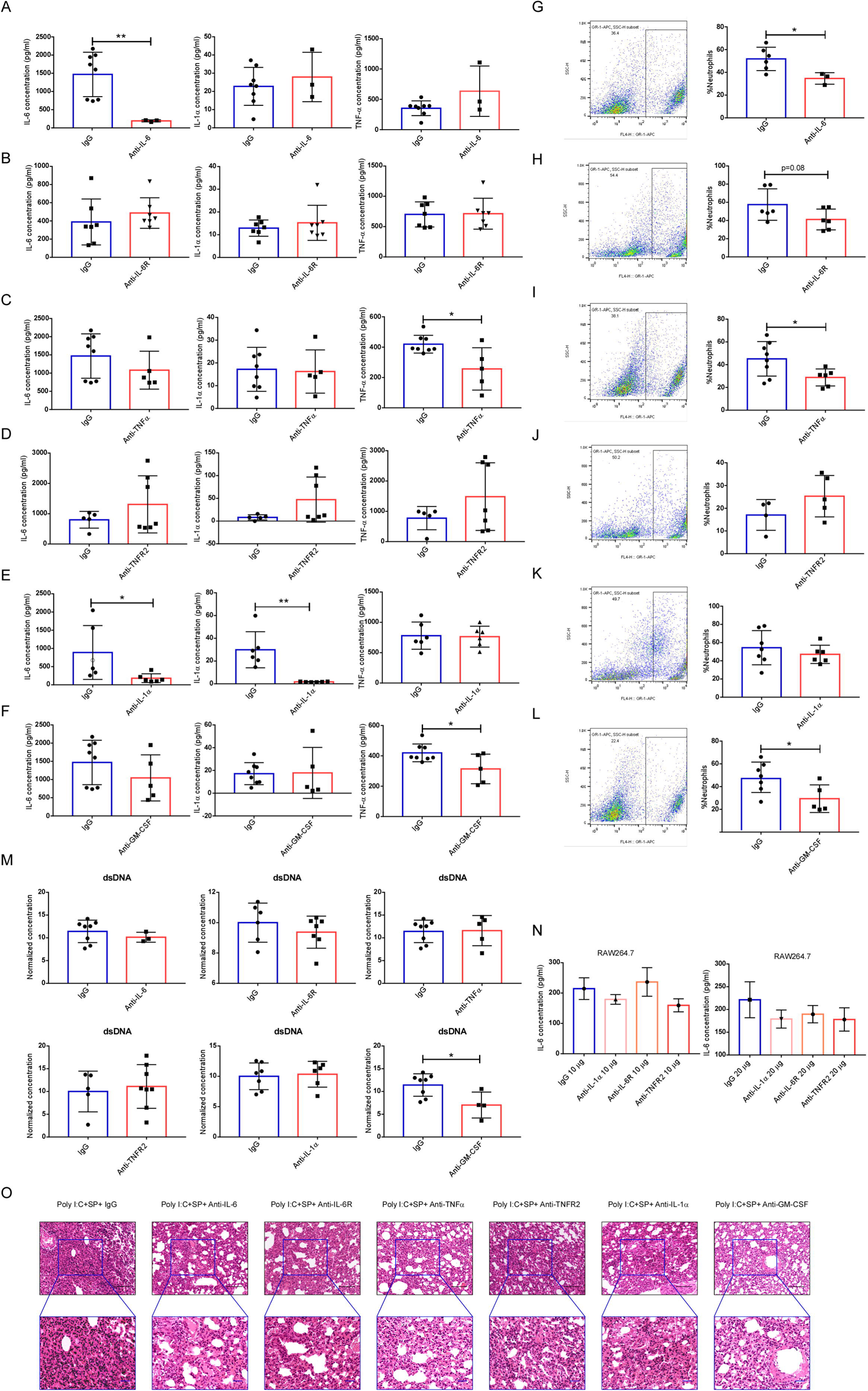
Blocking of inflammatory cytokines. BAL IL-6, IL-1α and TNFα concentrations from anti-IL-6 mAbs (a), anti-IL-6R mAbs (b), anti-TNFα mAbs (c), anti-TNFR2 mAbs (d), anti-IL-1α mAbs (e), anti-GM-CSF (f) mAbs treated mice. * *P* < 0.05, ** *P* < 0.01. Cellular composition in BAL, anti-IL-6 (g), anti-IL-6R (h), anti-TNFα (i), anti-TNFR2 (j), anti-IL-1α (k) or anti-GM-CSF (l) mAbs treated mice. (m) Levels of dsDNA in the BAL. Poly I:C stimulate the IL-6 production in mouse macrophages (n) IL-6 concentration from cell culture medium. (o) Histological characteristics of lung injury and interstitial pneumonia. Black scale bar = 100 μm, blue scale bar = 50 μm

Infiltration of neutrophils caused the neutrophilic alveolitis during SARS-CoV-2 challenge. The percent of neutrophils in BAL were tested after anti-IL-6 (Figure 3g), anti-IL6R (Figure 3h), anti-TNFα (Figure 3i), anti-TNFR2 (Figure 3j), anti-IL-1α (Figure 3k) and anti-GM-CSF (Figure 3l) treatment. The result indicated that neutralizing of cytokines IL-6, TNFα and GM-CSF or blocking the IL-6R could reduce the infiltration of neutrophils. Concentrations of dsDNA were no reduction after antibodies treatment (Figure 3m). Next, we treated RAW264.7 cells with Poly I:C and evaluated the production of IL-6 in the medium. The results indicated a reduced trend of IL-6 while treating with 20 μg antibodies although not significantly (Figure 3n). Our histological analysis results indicated that treatment with neutralizing or blocking mAbs of inflammatory cytokines could partially reduce the neutrophilic inflammation and interstitial edema in the lung, but not anti-TNFR2 (Figure 3o). Therefore, potent therapeutic efficacy in a synergistic manner can be considered in clinical.

NETs can also be generated by macrophages^13^, and strongly contribute to acute lung injury during virus infection^14-16^. We hypothesized that the SARS-CoV-2 mimic stimulates the macrophages and produce inflammatory cytokines, such as IL-1α, IL-6 and TNF-α, recruit and activate neutrophils. P38 and ERK are the signaling mediators of NETs formation. MPO and Elastase as the peroxidase enzyme in neutrophils and essential for NETs formation ^17,18^. TLR3/dsRNA complex inhibitor can block the interactions between TLR3 and Poly I:C. JAK inhibitor can block the downstream signaling of IL-6. In our study, we found that the inhibition of P38 reduces IL-1α and inhibition of JAK reduces IL-6 significantly (Figure 4a). We noticed a decrease in the infiltration of neutrophils after treatment of P38, ERK and MPO inhibitors, which all targeted activities of neutrophils (Figure 4b). There was no significant difference in the concentrations of dsDNA after the treatment of those inhibitors (Figure 4c). Furthermore, we performed the cell assay to validate the effect of TLR3/dsRNA inhibitor and JAK inhibitors on mouse primary macrophages. TLR3/dsRNA inhibitor did not reduce the IL-6 production at 45 μM. JAK1/2 inhibitor Baricitinib reduced the production of IL-6 significantly, but not Febratinib (JAK2 inhibitor) (Figure 4d). Histological analysis result indicated that inhibitors of P38, ERK, MPO and Baricitinib (JAK inhibitor) obviously decreased the neutrophilic inflammation and interstitial edema in the lung (Figure 4e).

**Figure 4:**
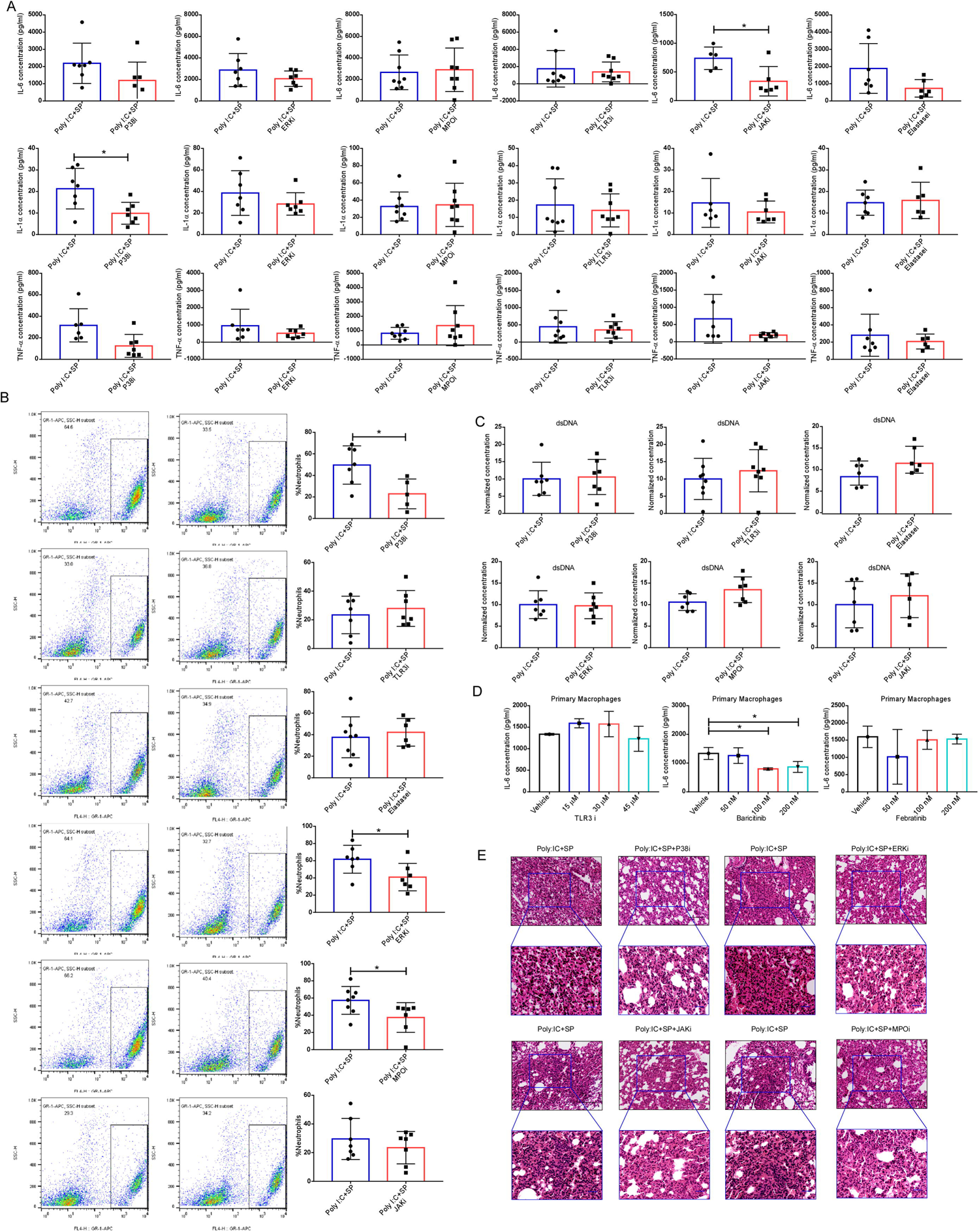
Blocking of inflammatory related signaling pathways. (a) BAL IL-6, IL-1α and TNFα concentrations. * *P* < 0.05. (b) Percent of neutrophils in BAL. * *P* < 0.05. (c) Levels of dsDNA in the BAL. Poly I:C stimulate the IL-6 production in mouse primary macrophages (d) IL-6 concentration from cell culture medium. * *P* < 0.05. (e) Histological characteristics of lung injury and interstitial pneumonia. Black scale bar = 100 μm, blue scale bar = 50 μm.

**Figure 5:**
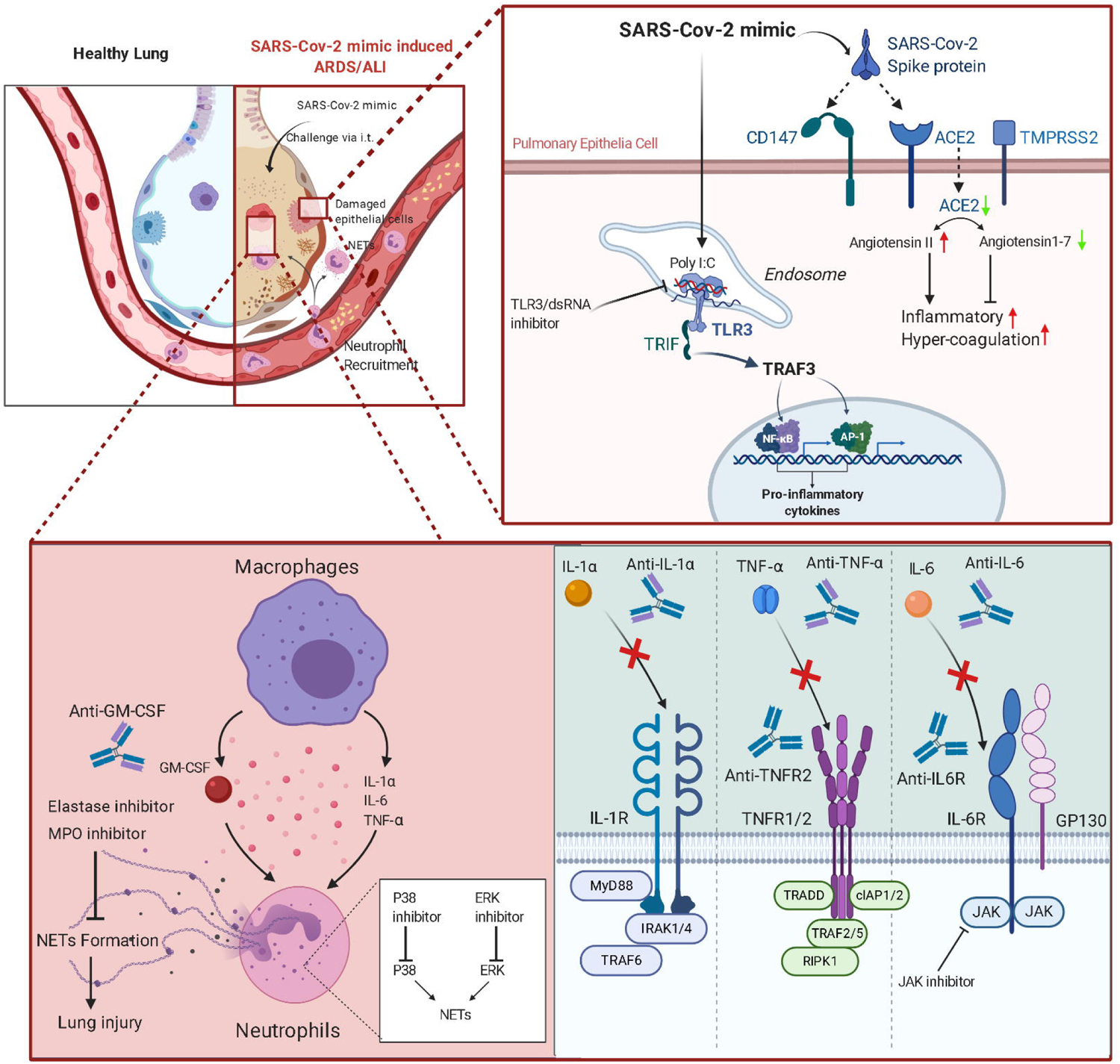
Graphical abstract of SARS-CoV-2 mimic and applications of therapeutic mAbs and inhibitors.

## Discussion

SARS-CoV-2 has posed a serious threat to global public health. Antiviral drugs, such as hydroxychloroquine, chloroquine and remdesivir did not achieve an ideal therapeutic effect^19,20^. In order to understand the molecular mechanism of SARS-CoV-2 induced ARDS/CSS and find an effective strategy to prevent and remedy this highly infectious disease, we have developed a murine ARDS model, to mimic pathological changes in COVID-19 patient. Coronavirus includes a large genomic RNA and can stimulate TLR3/7 upon infection^21-25^. We used Poly I:C to mimic the effect of viral RNA. Moreover, the unique outcome of recombinant SP was showed in a TNFα outbreak featured CSS when compared with Poly I:C challenged mice. The TNFα storm revealed that the underlying mechanism of CSS happened in COVID-19 patients, not just IL-6 or IL-1α, TNFα may play a more critical role. As a new target in our model, anti-TNFα may need a tight treatment window, the TNFα burst was observed after 6 h administration of SARS-Cov-2 mimic and disappeared at 24 h. Furthermore, IL-6 maintained its high level during the administration of SARS-CoV-2 mimics, anti-IL-6, or related signaling may provide a relatively broad treatment window.

Impaired inflammatory responses in severe COVID-19 patients provide a distinct pattern of COVID-19 progression in host immune response to the SARS-CoV-2 infection, with exhausted lymphocytes, high neutrophil to lymphocyte ratio, CSS in severe cases of COVID-19^9,26-29^. CSS was found to be the major cause of morbidity in patients with SARS-CoV and MERS-CoV^30^, with the presence of IL-6 in the plasma being a hallmark of severe infections. Clinical trials using IL-6 and IL-6R antagonists are underway^31^. In our research, we first tested the effect of IL-6 or IL-6R mAbs, IL-6 in BAL was affected, but IL-1α and TNFα no changes. Importantly, treatment of IL-6 or IL-6R mAbs reduced neutrophil infiltration. This result may indicate that blocking IL-6 signaling shows partially benefit from the SARS-CoV-2 mimics induced neutrophilic alveolitis, and CSS in lung did not appear to effective mitigation. Furthermore, blocking of JAK-STAT3 signaling also could inhibit the IL-6 mediated signaling transduction. For blocking TNFα signaling, we performed the TNFα and TNFR2 mAbs. TNF exerts its effect by stimulation of two different receptors, TNFR1 and TNFR2. TNFR1 is expressed in all cell types, but TNFR2 is only expressed in certain cell types, such as myeloid cells, glial cells and T and B cell subsets. Neutralizing TNFα via mAbs reduced the production of TNFα in our model, IL-6 and IL-1α not affected. Treatment of TNFα mAbs, but not TNFR2 mabs reduced neutrophil infiltration. Those results indicated that TNFα signaling shows partially reduced neutrophilic alveolitis and benefit from SARS-Cov-2 mimic induced CSS. GM-CSF is a cytokine found in high levels in patients with COVID-19, anti-GM-CSF mAbs was used for the treatment of COVID-19 clinical trial (NCT04400929)^2^. In our present data, anti-GM-CSF reduced neutrophil infiltration and decreased the TNFα during SARS-Cov-2 mimic challenge. Even we did not find GM-CSF enriched in BAL, but anti-GM-CSF treatment shows protection effect from neutrophilic alveolitis and NETs induced lung injury.

It was reported that macrophages in the lungs may contribute to inflammation by producing multiple cytokines/chemokines and recruiting more inflammatory monocytic cells and neutrophils ^32,33^. We have applied various signal pathway inhibitors to block the dysregulated macrophages and neutrophils. TLR3/dsRNA Complex inhibitor blocked the dsRNA and TLR3 interaction, but in our present data, TNFα in BAL cannot induced by single Poly I: C. and SARS-Cov-2 includes a single strand RNA genome, TLR7 also sense the viral RNA^34^.

To prevent sustained activation of neutrophils, we applied P38 inhibitor SB203580, ERK inhibitor Tauroursodeoxycholate and MPO inhibitor 4-Aminobenzoic hydrazide in the SARS-CoV-2 mimic model. It was reported that MPO is a functional and activation marker of neutrophils. Elastase was produced by neutrophils and can mediate lung injury. Inhibitors of MPO and elastase can directly reduce the acute lung injury at the inflammatory site^18,35,36^. In our study, inhibitors include P38 inhibitor, ERK inhibitor, MPO inhibitor and elastase inhibitor can reduce the neutrophil infiltration into lung, but not elastase. Here we used the neutralizing or blocking antibodies, essential pathway inhibitors of the innate and adaptive immune responses to target the most abundant cytokines, NETs and other pro-inflammatory factors to prevent the SARS-CoV-2 caused CSS and ARDS. Our data indicate that potential combination of those neutralizing/blocking antibodies or inhibitors may contribute to the treatment of CSS and ARDS, and should be investigated further.

Taken together, we established a non-infectious, high safety and time-saving murine robustness model by using Poly I:C and SP, which mimic the pathological changes of SARS-CoV-2 caused CSS and ARDS. This model can be available for research on COVID-19 caused CSS and ARDS, addressing the immunopathology alterations, and exploring new therapies of COVID-19.

## Methods and material

### Experimental Animals

Male BALB/c mice (8-10 weeks) free of pathogens were purchased from Beijing Vital River Laboratory Animal Technology Co., Ltd. (Beijing, China). Mice were housed in a pathogen-free environment under conditions of 20°C ± 2°C, 50% ± 10% relative humidity, 12-h light/dark cycles. They were provided with food and water *ad libitum.* All experimental procedures involving animals were approved by the Ethics Review Commission of Zhengzhou University (following internationally established guidelines).

### COVID-19 ARDS murine model and treatment

Mice were anesthetized via intraperitoneal (IP) injection with Pentobarbital Sodium (50 mg/kg). A small incision was made over the trachea, and the underlying muscle and glands were separated to expose the trachea. Mice were intratracheally administered with freshly mixed Poly I:C (Poly I:C-HMW, Invivogen, tlrl-pic) 2.5 mg/ml and SARS-CoV-2 recombinant spike protein (ECD-His-tag, Genescript, Z03481) 15 μg (in saline), followed by 100 μL air. 2.5 mg/kg Poly I:C, FC control (ACRO, P01857-1), 15 μg SARS-CoV-2 recombinant spike protein and saline were administered intratracheally independent at the same volume as control.

Blocking and neutralizing antibodies anti-IL-1α (InVivoMab, BE0243), IL-6R (InVivoMab, BE0047), and TNFR2 (InVivoMab, BE0247) were administrated intraperitoneally as a single dose of 200□μg 24□h in prior. TLR3/dsRNA inhibitor (Merck, 614310) was administrated via i.p. at 50 mg/kg 2 h prior to administration of SARS-CoV-2 mimics. MPO inhibitor (Merck, A41909) was administrated via i.p. at 50 mg/kg per day, 3 d prior to administration of SARS-CoV-2 mimics. P38 inhibitor (MCE, HY-10256) was administrated via i.p. at 20 mg/kg per day, 3 d prior to administration of SARS-CoV-2 mimics. ERK inhibitor (MCE, HY-19696A) was administrated via i.p. at 100 mg/kg per day, 3 d prior to installation of SARS-CoV-2 mimics. Elastase Inhibitor (GLPBIO, GC11981) was administrated via i.p. at 5 mg/kg per day, 3 d prior to administration of SARS-CoV-2 mimics. JAK inhibitor (MCE, HY-15315) was administrated via i.p. at 20 mg/kg per day, 3 d prior to administration of SARS-CoV-2 mimics.

### Analysis of BAL samples

6 h after the administration, mice were anesthetized via intraperitoneal (IP) injection of Pentobarbital Sodium 50 mg/kg. A 26G venous indwelling needle hose was inserted into the exposed tracheal lumen, and then the airway was washed three times with 1 ml saline each, the first lavage fluid sample was kept separately to test cytokines.

### Multiplex cytokines assay

BAL was collected from mice lung after anesthetized, the mice BAL were separated by centrifuging and stored at −80□. BAL sample collection was performed as previously described. Next, concentrations of mice inflammatory related cytokines IL-1α, IL-1β, IL-6, IL-10, IL-12p70, IL-17A, IL-23, IL-27, MCP-1, IFN-β, IFN-γ, TNF-α, and GM-CSF were measured by LEGENDplex™ Mouse Inflammation Panel (13-plex, Biolegend), the datas were harvest by flow cytometry using FACSCalibur (BD).

### Flow Cytometry analysis

Fluorescent labeled antibodies were performed to quantify neutrophils and macrophages. Unconjugated anti-mouse CD16/CD32 (Biolegend, 101320) was used for blocking Fc receptors, APC labeled anti-mouse Ly-6G/Ly-6C (Gr-1) and FITC labeled anti-mouse F4/80 were performed for neutrophils and macrophages, incubated 30 min on ice, protect from light. Washed three times by PBS and centrifuged to remove the supernatant and responded in 150 μL PBS. Samples were analyzed in the BD FACSCalibur (BD).

### Histopathology Analysis

An independent experiment was performed for the histopathology analysis of pulmonary. The mouse lungs were removed intact and weighted, then fixed in 10% formalin and paraffin-embedded. Three different fields from a lung section were evaluated, 3 μm sections were sliced on a Leica model rotary microtome and stained with hematoxylin-eosin. Histological analysis was subjected by two independent skilled pathologists, in double-blind.

### Mouse macrophage cell culture

RAW264.7 cell was cultured in DMEM medium supplemented with penicillin (100 units/mL), streptomycin (100 μg/mL), and 10% FBS (Biological Industries, Kibbutz Beit-Haemek, Israel).

### Compounds treated to RAW264.7

RAW264.7 cells (6×10^5^ cells/well) were seeded into 12-well plates and cultured for 24 h in incubator. The cells were treated with a series dilution of compound X; meanwhile, Poly I:C (10 μg/ml) were added into medium in a 12-well plate. Meanwhile, RAW264.7 cell was subjected to DMEM with or without Poly I:C as a positive or negative control. The cells were maintained at 37°C in a 5% CO_2_ incubator for 12 h, harvested the supernatant for the detection of IL-6 secretion.

### Mouse primary macrophage cell culture

Bone were gently separated from mouse, and the connecting muscles and soft-tissues were all removed from the bone. The macrophages contain in the bone were washed out by DPBS using 1 ml syringe. Suspended the eluate and centrifuged at 1000 rpm for 5 min. Washed the pellets by DPBS and resuspended the cell pellets by RPMI-1640 containing penicillin(100 units/mL), streptomycin (100 μg/mL), 10% FBS (Biological Industries, Kibbutz Beit-Haemek, Israel) and 50 ng/ml M-CSF (peprotech), plated the cells in a low attachment culture dish and maintained at a 5% CO_2_ incubator. Renew the medium of fresh RPMI-1640 complete medium contained 50 ng/ml M-CSF every other day until the density of adherent cells reached 90%. Digested the cells and plated 6×10^5^ cells/well into 12-well plates, cultured for 24 h and treated the Poly I:C and compound to the cell using blank RPMI-1640 medium. This method is similar to the treatment of RAW264.7.

### Dissociated of primary lung cells mix

Took out the whole lung tissues of the mice and washed by pre-cold PBS containing DNase I (0.01 mg/ml) (Sigma), penicillin (100 units/mL), and streptomycin (100 μg/mL). Tattered the lung tissues into small pieces in a sterile centrifuge tube by scissors, suspended the small pieces by 1 mg/ml Type I collagenase (Sigma) supplied by 0.01 mg/ml DNase I and digested in a 37□ shaking incubator for 30 min. Filtered the solution with a 200 mesh sieve and centrifuged at 1000 rpm for 5 min, subsequently. After washed the pellets twice by pre-cold PBS containing DNase I (0.01 mg/ml), resuspended the pellets by RPMI-1640 containing 10% FBS (Biological Industries, Kibbutz Beit-Haemek, Israel) and seeded into 12-well plate. Harvested the supernatant after 24 h culture and removed the floating cells by centrifuge at 3000 rpm for 5 min. The ELISA assay was performed to detect the IL-6 concentration following the manufacturer’s instructions.

### Statistical Analysis

Statistical analysis was performed using GraphPad Prism 7.0 (GraphPad Software, United States). Specific statistical methods and comparisons made by methods as described in figure legend. Comparison between two groups were performed by paired Student’s t-test or unpaired Student’s t-test. *P* < 0.05 was regarded as statistically significant and marked with a star, data was reported as mean□±□SEM, and error bars indicate SEM.

## Graphical Illustrations

Schematic illustrations were established with BioRender (BioRender.com).

## Author contributions

Z.G.D., T.X.G. and S.M.Z. designed the study. T.X.G., S.M.Z., G.G.J, M.Q.S., Y.F.Z., R.Z., F.Y. M., Y.Q. Z., K.K.W. performed the main experiment. H.L., M.X.X, W.H. collected and analyzed the raw data. Z.G.D., T.X.G. and S.M.Z. wrote the manuscript. X.L., C.D.D., K.D.L. revised the manuscript. All authors read and approved the final manuscript

